# Evaluating the breadth of neutralizing antibody responses elicited by infectious bursal disease virus (IBDV) genogroup A1 strains using a novel chicken B-cell rescue system and neutralization assay

**DOI:** 10.1101/2022.06.03.494759

**Authors:** Vishwanatha R. A. P. Reddy, Salik Nazki, Andrew J. Brodrick, Amin Asfor, Joanna Urbaniec, Yasmin Morris, Andrew J. Broadbent

## Abstract

Eight infectious bursal disease virus (IBDV) genogroups have been identified based on the sequence of the capsid hypervariable region (HVR) (A1-8), yet many vaccines are based on A1 strains. Given reported vaccine failures, there is a need to evaluate the ability of vaccines to neutralize the different genogroups. To address this, we used a reverse genetics system and the chicken B-cell line DT40 to rescue a panel of chimeric IBDVs and perform neutralization assays. Chimeric viruses had the backbone of a lab-adapted strain (PBG98) and the HVRs from diverse field strains: classical F52-70 (A1), US-variant Del-E (A2), Chinese-variant SHG19 (A2), very-virulent UK661 (A3), M04/09 distinct (A4), Italian ITA-04 (A6), and Australian-variant Vic-01/94 (A8). Rescued viruses showed no substitutions at amino-acid positions 253, 284, or 330, previously associated with cell-culture adaptation. Sera from chickens inoculated with wt (F52-70) or vaccine (228E) A1 strains had the highest mean virus neutralization (VN) titers against the A1 virus (log_2_ 15.41 and 12.66), and the lowest against A2 viruses (log_2_ 7.41-7.91, p=0.0001-0.0274), consistent with A1 viruses being most antigenically distant from A2 strains, which correlated with the extent of differences in the predicted HVR structure. VN titers against the other genogroups ranged from log_2_ 9.32-13.32, and A1 strains were likely more closely antigenically related to genogroups A3 and A4 than A6 and A8. Our data are consistent with field observations, validating our method, which can used to screen future vaccine candidates for breadth of neutralizing antibodies, and evaluate the antigenic relatedness of different genogroups.

**Importance:** There is a need to evaluate the ability of vaccines to neutralize diverse IBDV genogroups, and to better understand the relationship between HVR sequence, structure, and antigenicity. Here, we used a chicken B cell-line to rescue a panel of chimeric IBDVs with the HVR from seven diverse IBDV field strains, and conduct neutralization assays and protein modelling. We evaluated the ability of serum from vaccinated or infected birds to neutralize the different genogroups. Our novel chicken B-cell rescue system and neutralization assay can be used to screen IBDV vaccine candidates, platforms, and regimens for the breadth of neutralizing antibody responses elicited, evaluate the antigenic relatedness of diverse IBDV strains, and when coupled with structural modelling, elucidate immunodominant and conserved epitopes to strategically design novel IBDV vaccines in the future.

## Introduction

Infectious bursal disease virus (IBDV), a member of the genus *Avibirnavirus* in the family *Birnaviridae*, is a highly contagious and immunosuppressive virus that infects commercial poultry worldwide, and is ranked among top five infectious problems of chickens (1, 2). IBDV is a non-enveloped virus with a bi-segmented double-stranded RNA genome comprised of segment A (3.2 Kb) and segment B (2.8 Kb), enclosed within an icosahedral capsid. Segment A has two partially overlapped open reading frames (ORF), where ORF A1 encodes the non-structural viral protein VP5 that is reported to be involved in virus egress (3), and ORF A2 encodes a large polyprotein that undergoes cleavage by the protease VP4 to yield VP2, VP4, and VP3 (4). VP2 is the capsid protein, and VP3 is a multifunctional protein that binds the dsRNA genome and may help form a scaffold between the genome and the capsid (5, 6). Segment B has one ORF that encodes the RNA dependent RNA polymerase (VP1) enzyme, which is involved in viral genome replication (7). Both segment A and B contribute to the pathogenicity of IBDV (8).

The VP2 capsid is known to be an important immunodominant protein of IBDV and is the major target of neutralizing antibodies, which are thought to be the main correlate of protection. Within VP2, there is a so-called “hypervariable region” (HVR), located between amino acids 220 to 330, which is subject to the most intense immune selection pressure and antigenic drift. IBDV strains have been divided into eight genogroups based on the sequence diversity of the HVR, termed genogroups A1-A8 (9, 10). Furthermore, within the HVR, there are four hydrophilic loops of amino acids that project out from the tip of the VP2 molecule. These loops are termed P_BC_, P_DE_, P_FG_, and P_HI_, and are reported to contribute to IBDV pathogenicity and antigenicity (11–14).

Recently, there has been an increase in reports of IBDV vaccine failures throughout the globe, which has been attributed to the emergence of variant IBDV strains containing mutations in the HVR (9, 15–17). However, how IBDV HVR sequence diversity relates to antigenic diversity is poorly understood, and there is a need to conduct fundamental research to provide new information on how sequence changes in the HVR relate to changes in antigenicity, and identify immunodominant epitopes. In addition, there is an urgent need to conduct applied research to evaluate the breadth of neutralising antibodies elicited by commercial IBDV vaccines, to evaluate their use in different geographical regions, against different genogroups, and to determine the potential for immune escape. However, until now, conducting these studies has been difficult because field strains of IBDV have a preferred tropism for B cells, and do not replicate well in immortalised adherent cell-lines, without prior adaptation associated with mutations in the HVR that could change antigenicity and virulence (12–14). As such, field strains are typically propagated by passage *in vivo*, by harvesting the bursa of Fabricius (BF) from infected birds, or *in ovo*, by inoculating embryonated eggs (18–20). Moreover, the ability to rescue a molecular clone of IBDV was, until recently, limited to laboratory strains of IBDV that were adapted to replicate within chicken embryo fibroblasts (CEFs), DF-1, QM7 or Vero cells, further hampering the ability to study how individual mutations within the HVR of field strains contribute to antigenicity. Recently, we and others demonstrated that field strains of IBDV can replicate within primary chicken bursal cells and the immortalised chicken B-cell line DT40 (19, 21–25). Moreover, primary chicken bursal cells were used to rescue a molecular clone of a field strain of IBDV for the first time in 2020 (26), thus enabling the ability to study how mutations in the IBDV HVR contribute to antigenicity and immune escape in field strains.

The sequences of the HVRs from diverse strains of IBDV are available in GenBank, but often the whole VP2 sequence is lacking. Taking advantage of the available HVR sequences, and our inhouse IBDV reverse genetics system (27), here we describe the rescue of a panel of seven chimeric IBDVs containing the HVR from diverse strains belonging to six different genogroups from different geographical regions, in the backbone of strain PBG98. The chimeric viruses were rescued in the chicken B cell line DT40, and subsequently used to determine the breadth of neutralising antibody responses elicited by virulent and vaccine strains belonging to genogroup A1.

## Materials and Methods

### Cell lines and antibodies

The chicken B-cell lymphoma cell-line, DT40 (ATCC cat number), was maintained in RPMI media supplemented with l-glutamine and sodium bicarbonate (Sigma-Aldrich), 10% heat-inactivated fetal bovine serum (FBS) (Sigma-Aldrich), trypots phosphate broth (Sigma-Aldrich), sodium pyruvate (Sigma-Aldrich) and 50 mM beta-mercaptoethanol (Gibco) (complete DT40 media) (28). The primary antibodies used in this study were raised against VP3 (27). In all immunofluorescent staining, primary antibodies were diluted 1:100, and secondary antibodies conjugated to Alexa 568 (Invitrogen, Thermo Fisher Scientific) were diluted 1:500 in a solution of bovine serum albumin (BSA; Sigma-Aldrich).

### Viruses

The virulent IBDV field strain F52/70 (29), and the very virulent (vv) IBDV field strain UK661 (30), were kind gifts from Dr Nicolas Eterradossi (ANSES, Ploufragen, France). Viruses were propagated *in vivo* by harvesting the bursa of Fabricius (BF) from experimentally inoculated chickens at 72 hours post infection (hpi). The bursal material was pooled from six chickens, and homogenized in Vertrel XF (Sigma-Aldrich, Merck), which separated into two phases. The upper phase was harvested and layered on top of a 30% sucrose solution and ultra-centrifuged at 27,000rpm. The resulting pellet was resuspended in PBS. The lyophilized live attenuated vaccine, Nobilis strain 228E^®^ was obtained from Intervet (International BV, Boxmeer, Holland), and reconstituted as per the manufacturer’s instructions and titrated in 10 days old embryonated eggs.

### Titration of IBDV in DT40 cells

IBDV was diluted ten-fold in complete DT40 media in U-bottom 96-well plates (Falcon, Corning, UK), in quadruplicate, and DT40 cells were then added to diluted virus at 1 x 10^5^ cells/well. Cells were incubated in the presence of diluted virus for 3 days, fixed in 4% paraformaldehyde solution (Sigma-Aldrich) for 20 min, permeabilized with a solution of 0.1% Triton X-100 (Sigma-Aldrich) for 10 min, and blocked with a 4% BSA solution for 60 min. The cells were then incubated with a primary mouse monoclonal antibody raised against the IBDV VP3 protein (31) for 1 h at room temperature. Cells were washed with phosphate-buffered saline (PBS) and incubated with a goat-anti-mouse secondary antibody conjugated to Alexa fluor 488 or 568 (Thermo Fisher Scientific) for 1 h at room temperature in the dark. The cells were again washed and incubated for 10 min in a solution of 4’,6’-diamidino-2-phenylindole (DAPI) (Invitrogen, Thermo Fisher Scientific). Cells were imaged using a Leica DM IRB epifluorescence microscope. The highest dilution of the virus where 50% of the wells had a VP3 signal was considered as the end point, and the virus titer was determined from the tissue culture infectious dose-50 (TCID_50_), according to the method of Reed and Muench, and expressed as TCID_50_/mL (32).

### Titration of IBDV in embryonated hens’ eggs

IBDV was diluted ten-fold in PBS and inoculated onto the chorioallantoic membrane (CAM) of specific pathogen free (SPF) embryonated eggs at 10 embryonic days of age (ED10) and titrated as previously described (19). Briefly, inoculated eggs were incubated for 7 days at 37°C, whereupon embryos were humanely culled and observed for signs of pathology caused by the virus. The highest dilution of the virus where 50% of the embryos had IBDV-mediated pathology was considered as the end point, and the virus titer was determined from the egg infectious dose-50 (EID_50_) according to the method of Reed and Muench, and expressed as EID_50_/mL.

### Rescue of a molecular clone of IBDV in DT40 cells by electroporation

Reverse genetics plasmids encoding segments A and B from IBDV strain PBG98 (pPBG98A and pPBG98B) were constructed as previously described (27). DT40 cells of 1 × 10^7^ were resuspended in 100 μL Opti-MEM medium, and 10 μg of pPBG98A and pPBG98B were mixed with the cells. The mixture was then electroporated at 225 V and a pulse width of 2 ms of poring pulse. Forty eight hours post-electroporation (hpe), cell cultures were ‘fed” with fresh DT40 cells. Cultures continued to be fed every 72 hours, where fresh cells were added to old cells in a 3:1 ratio.

### Rescue of a panel of chimeric recombinant IBDVs with the backbone of PBG98 and the HVR of diverse field strains

The HVR sequences from seven diverse field strains were retrieved from GenBank database as follows: classical strain F52-70 (genogroup A1), accession number AY321953, US-variant strain Delaware-E (Del-E, genogroup A2), accession number AF133904, Chinese-variant strain SHG19 (genogroup A2), accession number MH879092, vv strain UK661 (genogroup A3), accession number NC_004178, M04/09 distinct strain (genogroup A4), accession number KM659895, Italian ITA-04 strain (genogroup A6), accession number JN852988, and Australian-variant Vic-01/94 strain (genogroup A8), accession number AF148076. For every strain, the HVR was comprised of 333 nucleotides that encoded 111 amino acids, numbered from residue 220 to 330. Seven plasmids encoding IBDV segment A were designed, each containing the HVR from a different field strain, and the rest of the segment from strain PBG98. Plasmids were synthesised by GeneArt (Thermo Fisher Scientific, UK) and cloned into a pSF-CAG-KAN vector (Addgene, UK) using restriction enzyme pairs Kpn1/ Nhe1. The resulting chimeric plasmids pPBG98/A/HVR-F52-70, Del-E, SHG19, UK661, M04/09, ITA-04 and Vic-01/94 were then sequenced using pSF-CAG-KAN vector forward primer 5’-CTACCATCCACTCGACACACC-3’ and reverse primer 5’-GTTGTGGTTTGTCCAAACTCATCA-3’ (Integrated DNA Technologies, Belgium). DT40 cells of 1 × 10^7^ were suspended in 100 μL Opti-MEM medium, and 10 μg of pPBG98/B and 10 μg of one of the pPBG98/A/HVR plasmids were added to the cells. The mixture was then electroporated at 225 V and a pulse width of 2 ms of poring pulse. Forty eight hpe, cell cultures were fed with fresh DT40 cells (one “passage”). Cultures continued to be fed every 72 hours, where fresh cells were added to old cells in a 3:1 ratio. Viruses were passaged no more than 5 times. The sequences of HVRs of the rescued chimeric viruses were confirmed by using forward primer 5’-GCCCAGAGTCTACACCAT-3’ and reverse primer 5’-ATGGCTCCTGGGTCAAATCG-3’ (Integrated DNA Technologies, Belgium) (10).

### Growth curves of chimeric recombinant IBDVs

DT40 cells were seeded into 24-well plates at a density of 1 × 10^6^ cells per well in triplicate for each time point. The next day, cells were infected with one of the seven recombinant chimeric viruses, or PBG98 recombinant and wild type viruses at an MOI of 0.0005 for 1 hour at 37°C, 5% CO_2_. The cells were washed and resuspended in complete DT40 media and incubated at 37°C, 5% CO_2_. The cell supernatant was collected at 12, 24, and 48 hours post infection (hpi) and the virus titer determined by titration onto additional DT40 cells. The TCID_50_ was calculated according to the method of Reed and Muench (32).

### Bioinformatics analysis of VP2 HVR

Multiple-sequence alignments were performed using MEGA 6. 06 of the HVR sequences obtained from GenBank, the sequences of the plasmids, and the sequences of the rescued viruses, and the translated amino acid sequences were compared, respectively. Amino acid identities of the HVR sequences were determined using the p-distance model.

### Structural modelling of chimeric VP2 molecules

The sequences of the chimeric VP2 genes we designed were translated *in silico* using SnapGene (version 6.0.2, GSL Biotech), and the amino acid sequences were modelled using a modified version of AlphaFold v2.1.0 (33). The models were then downloaded and processed using PyMol (version 2.5, Schrödinger) to isolate the HVR and highlight residues that differed from PBG98. The same modelling process was employed to predict the structures of the rescued viruses, with slight modification: The 333 nucleotide sequence of each HVR obtained by sequencing the rescued virus were translated with SnapGene, and a Python script was employed to generate “virtual” full-length VP2 chimeras, by replacing HVR residues 220-330 of the canonical PBG98 sequence with the residues determined by translation of the rescued virus sequences.

### Collection of serum from F52-70 and 228E infected chickens

Nine three-week-old specific pathogen free (SPF) chickens of the Rhode Island Red (RIR) breed were hatched and reared at The Pirbright Institute, randomly designated into the following groups: mock-inoculated with PBS (n = 3), inoculated with the virulent classical field strain, F52-70 (n = 3) and vaccinated with the IBDV live vaccine 228E (n = 3). Briefly, each bird was inoculated with 10^5^ TCID_50_ dose virus intranasally, in a total of 100μLof PBS; 50μL per nares. All animal procedures conformed to the United Kingdom Animal (Scientific Procedures) Act (ASPA) 1986, under Home Office Establishment, Personal and Project licenses, following approval of the internal Animal Welfare and Ethic Review Board (AWERB) at The Pirbright Institute.

### Quantification of anti-IBDV serum neutralizing antibody titers

Serum samples were heated at 56 °C for 30 minutes to inactivate complement factors and serially diluted two-fold from 1:20 to 1:40960. Diluted serum was incubated with 100 TCID_50_ of each of the 7 chimeric strains of IBDV for one hour at 37 °C, and the mixtures were incubated with 1 × 10^8^ DT40 cells in 96 well U-bottom plates. Four days post-inoculation, cells were fixed and stained with an anti-IBDV VP3 antibody and a goat-anti-mouse secondary antibody conjugated to Alexafluor 488 or 568. Wells were scored as either positive or negative for IBDV antigen by immunofluorescence microscopy, and the virus neutralization (VN) titer was expressed as log_2_ of the highest dilution where no VP3-positive cells were observed. Following scoring the wells, the fixed and stained cells were diluted in FACS buffer and the percentage of VP3-positive cells quantified for each well by flow cytometry.

### Statistical Analysis

Viral titrations, growth curves and antibody virus neutralization titers were analysed by one-way analysis of variance (ANOVA) with Tukey post hoc comparisons using GraphPad Prism version 7.01 (GraphPad Software, Inc., San Diego, CA). Results were considered significantly different when P < 0.05. Unless otherwise stated, the results were shown as mean ± standard deviation (SD).

## Results

### DT40 cells can be used to quantify the titer of IBDV, and anti-IBDV serum neutralising antibodies

The very virulent (vv) IBDV field strain UK661 was serially diluted ten-fold and the diluted viral stocks were frozen at −80 °C. Each diluted stock was subsequently thawed and subject to titration by TCID_50_ in DT40 cells, and by EID_50_ in embyronated chicken eggs. A linear regression analysis revealed that there was a significant linear relationship between the log_10_ TCID_50_ and the log_10_ EID_50_ as R^2^ = 0.9313 (Figure 1A), demonstrating that the titer of the IBDV field strain could be quantified by TCID_50_ using DT40 cells. Hyperimmune serum from birds inoculated with the IBDV vaccine strain 2512 was obtained from Charles River (Massachusetts, USA), serially diluted, mixed with the UK661 virus, and added to the DT40 cells. Three days post-inoculation, the percentage of IBDV-positive cells in each well was determined by flow cytometry. Positive cells were detected when the virus was mixed with the hyperimmune serum at a dilution greater than 1:51,200, but not when the serum was diluted to 1:25,600 or less (Figure 1B), demonstrating the proof of concept that neutralising antibody titers against IBDV field strains could be quantified using the DT40 cells. In negative controls, the UK661 virus was mixed with chicken serum that lacked anti-IBDV antibodies. IBDV positive cells continued to be detected at all concentrations of negative serum tested, demonstrating that no virus neutralization occurred (Figure 1C).

**Figure 1.**
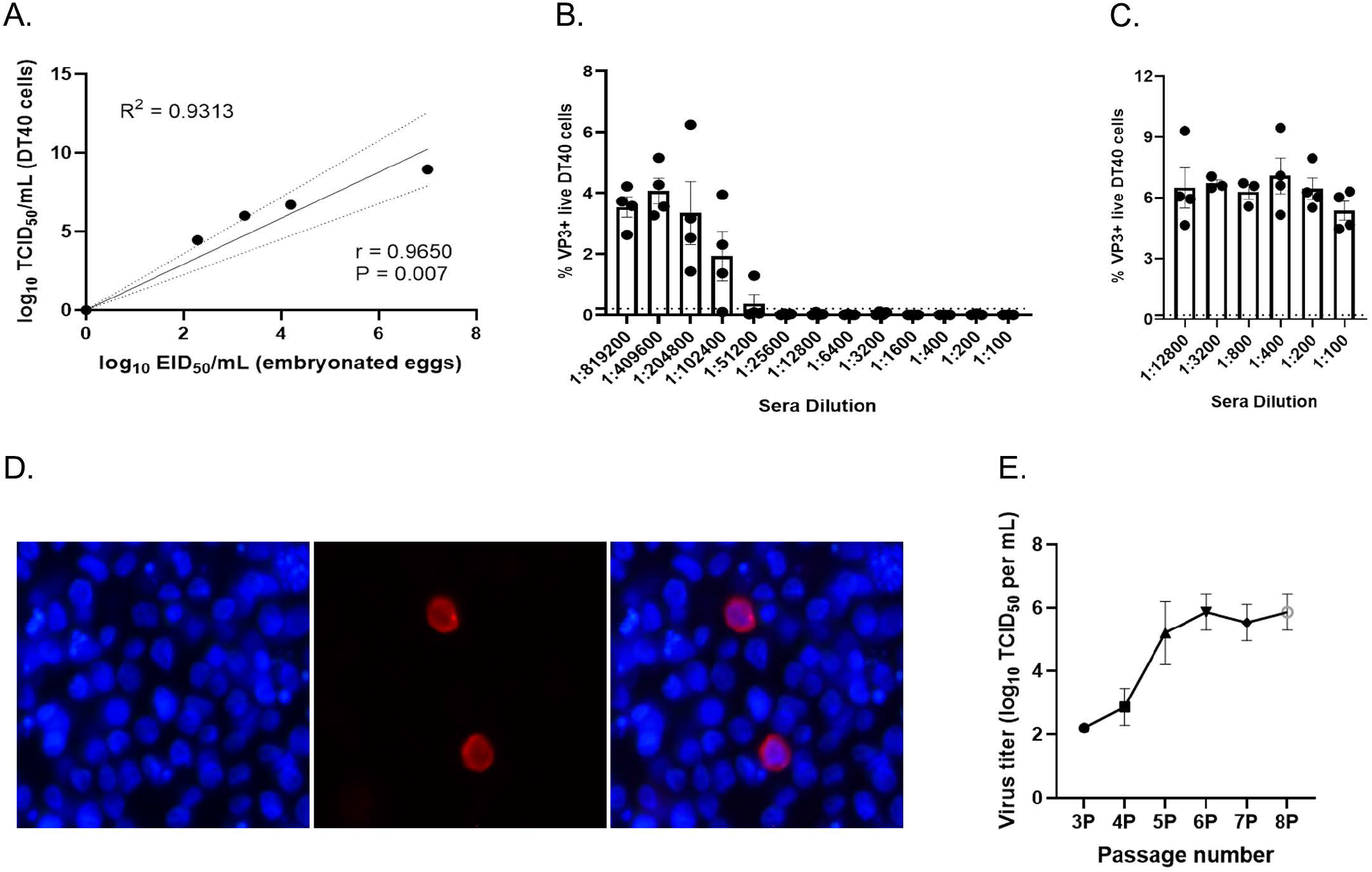
DT40 cells can be used to quantify the titer of IBDV, quantify the titer of anti-IBDV serum neutralising antibodies, and rescue a molecular clone of IBDV. The vv IBDV field strain UK661 was serially diluted ten-fold from 1:100 to 1:100,000, and the diluted stocks were frozen at −80 °C. Each diluted stock was subsequently thawed and subject to titration by TCID_50_ in DT40 cells, and by EID_50_ in embyronated chicken eggs, and a linear regression analysis was performed (A). Hyperimmune serum from birds inoculated with the IBDV vaccine strain 2512 (genogroup A1) was obtained from Charles River. The serum was heat inactivated, serially diluted two-fold, and mixed with 100 TCID_50_ of UK661. The mixture was added to DT40 cells in quadruplicate, and after 3 days, wells were fixed and stained with an antibody against the IBDV VP3 protein and a secondary antibody conjugated to a fluorophore. The wells were either scored positive or negative for the presence of IBDV antigen by immunofluorescence microscopy, and the percentage of positive cells in each well was quantified by flow cytometry to calculate the titer of the neutralising antibodies in the serum (B), compared to control serum from SPF birds that did not contain antibodies against IBDV (C). Each point represents the % of VP3+ DT40 cells in one well, the bar represents the mean, and error bars represent the standard deviation of the mean. The horizontal dashed line represents the limit of detection by TCID_50_ (B and C). Plasmids pBG98A and pPBG98B were electroporated to DT40 cells, and cell cultures were fed with fresh DT40 cells every 72 hours. Cells were fixed and stained with an antibody against IBDV VP3, a secondary antibody conjugated to AlexaFluor 568, and the nuclei were stained with DAPI. The IBDV antigen was present in the cytoplasm of the DT40 cells (VP3 viral antigen in red, DAPI in blue) (D). At each passage, the supernatant of the cultures was harvested, and serially diluted 10-fold in additional DT40 cells, to determine the titer as described by Reed & Muench. Three biological repeats were titrated and the mean titer plotted for each passage. Error bars represent the standard deviation of the mean (E).

### DT40 cells can be used to rescue a molecular clone of IBDV

DT40 cells were electroporated with reverse genetics plasmids pPBG98A and pPBG98B to rescue a molecular clone of IBDV strain PBG98 that was passaged by feeding the cultures with fresh DT40 cells. Rescue of the recombinant PBG98 virus was confirmed by epifluorescence microscopy following staining with an anti-VP3 antibody (Figure 1D). Viral titers increased steadily from passage three up to passage six, after which replication reached a plateau (Figure 1E). These data demonstrated the proof of concept that electroporation of DT40 cells was a successful method to rescue recombinant IBDV. Taken together, our experiments demonstrated that DT40 cells could be used to rescue recombinant IBDV strains, titrate field strains, and perform neutralization assays, and therefore made a suitable system for evaluating IBDV antigenicity.

### Rescue of chimeric IBDVs containing the HVR from diverse IBDV strains

Seven chimeric, recombinant IBDVs were rescued in DT40 cells, each with the backbone of the PBG98 strain and the VP2 HVR from a different field strain from a different geographical location, spanning six of the eight known genogroups based on segment A (Figure 2A). The field strains were classical strain F52-70 (genogroup A1), US-variant strain Delaware-E, Del-E (A2), Chinese-variant strain SHG19 (A2), vv strain UK661 (A3), M04/09 distinct strain (A4), Italian ITA-04 strain (A6), and Australian-variant Vic-01/94 strain (A8), and the HVR sequences were obtained from GenBank (Figure 2B). Virus rescue was confirmed by immunofluorescence microscopy. There was no significant difference in the peak titers between the chimeric strains, or between the recombinant and wild-type PBG98 strain (Figure 2C), demonstrating that the replication kinetics of all seven rescued chimeric viruses were similar, irrespective of the sequence of the HVR.

**Figure 2.**
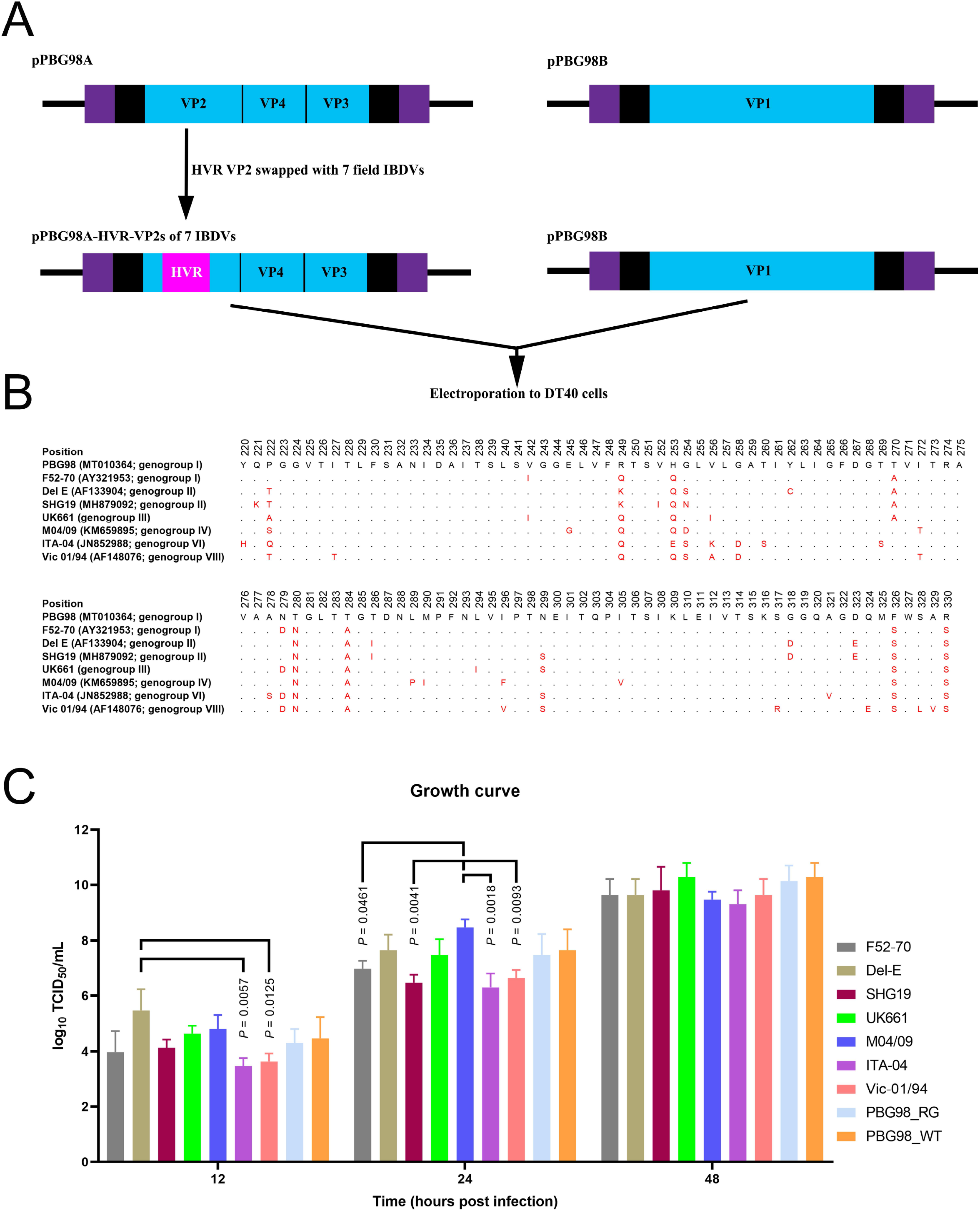
Construction of a panel of chimeric IBDVs with the backbone of the lab adapted PBG98 strain and the HVR from diverse field strains. Reverse genetics plasmids encoding segment A of the lab-adapted strain PBG98 (pPBG98A) were designed where the 333 nucleotides that encode the 111 amino acids (residues 220 to 330) of the HVR were swapped for one of seven field strains (pPBG98A-HVR-VP2s of 7 IBDVs). The plasmids were co-electroporated with the reverse genetics plasmid encoding segment B (pPBG98B) into DT40 cells to rescue the viruses (A). The HVR amino acid sequences from the seven field strains F52-70, Del-E, SHG19, UK661, M04/09, ITA-04 and Vic-01/94 were aligned with PBG98. The accession numbers and genogroup numbers are given in parenthesis. Conserved residues are depicted as black dots and different residues are highlighted in red (B). The replication kinetics of the seven recombinant chimeric IBDVs was determined in triplicate by titration of infected cell supernatants at 12, 24 and 48 hpi in DT40 cells, expressed as log_10_ TCID_50_/mL, and the mean plotted (error bars represent standard deviation of the mean) (C).

### Analysis of the HVR sequences of the rescued viruses

The sequences of the HVRs in the reverse genetics plasmids, and the rescued viruses that were passaged in DT40 cells, were compared to the sequences in GenBank (Figure 3). The sequences of the HVRs in the plasmids were identical to the corresponding GenBank sequences. Moreover, while it is known that IBDV adaptation to adherent-cell culture is mediated by amino acid changes at positions 253, 279, 284 and 330 (12, 13, 26, 34), we observed no change in the HVR at these positions in the majority of the chimeric IBDVs rescued in DT40 cells, compared to the GenBank sequences, except for amino acid position 279, which had an asparagine (N) to histidine (H) mutation (N279H) in strains Del-E and M04/09 (Figure 3). We did, however, note the following amino acid changes in the HVRs of the passaged viruses: S251I and S315Y (F52-70), C262Y and N279H (Del-E), T250S and S251I (UK661), V256L and N279H (M04/09), and T227I, T272I, S299N, E324Q, L328S, V329A (Vic 01/94).

**Figure 3.**
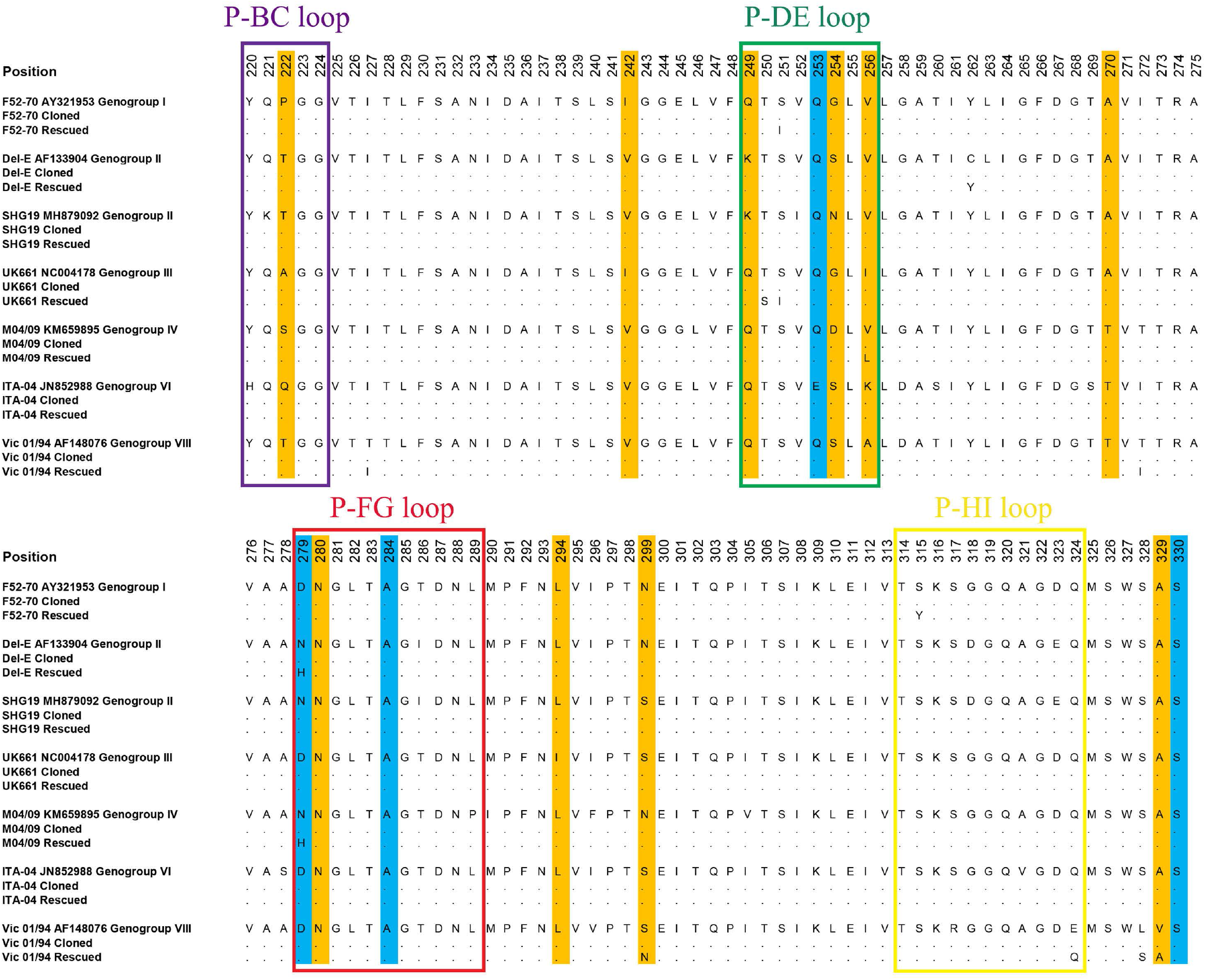
Sequencing analysis of the plasmids and viruses. The nucleotide sequences of the HVRs encoded by the seven plasmids, and present in the seven rescued and DT40-passaged viruses, were compared to the sequences in GenBank (Accession numbers provided) for strains F52-70, Del-E, SHG19, UK661, M04/09, ITA-04 and Vic-01/94. For each strain, the sequence in GenBank is displayed and conserved residues in the plasmid and the rescued virus are depicted as black dots, and different residues were listed. The four hydrophilic loops (P-BC, P-DE, P-FG, and P-HI), important for antigenicity, are boxed. Mutations previously reported to be involved in the adaptation of IBDV to adherent cell culture (positions 253, 279, 284 and 330) are highlighted in blue, and other common variable positions are shaded in orange.

### Analysis of the HVR structures

The structure of the chimeric VP2 molecules was predicted by AlphaFold and compared to the predicted structure of the genogroup A1 strain PBG98 (Figure 4). Structural modelling revealed that viruses belonging to genogroup A2 (Del-E and SHG19) had more extensive changes on the axial view of the VP2 molecule compared to PBG98 than the other genogroups (highlighted in orange) (Figure 4A). Interestingly, when the predicted structure of the chimeric VP2 molecules from the rescued and DT40-cell passaged viruses was compared to the predicted structure from the GenBank sequences for the corresponding strain, the majority of amino acid mutations associated with DT40 cell passage (highlighted in purple) were not on the axial view (Figure 4B).

**Figure 4.**
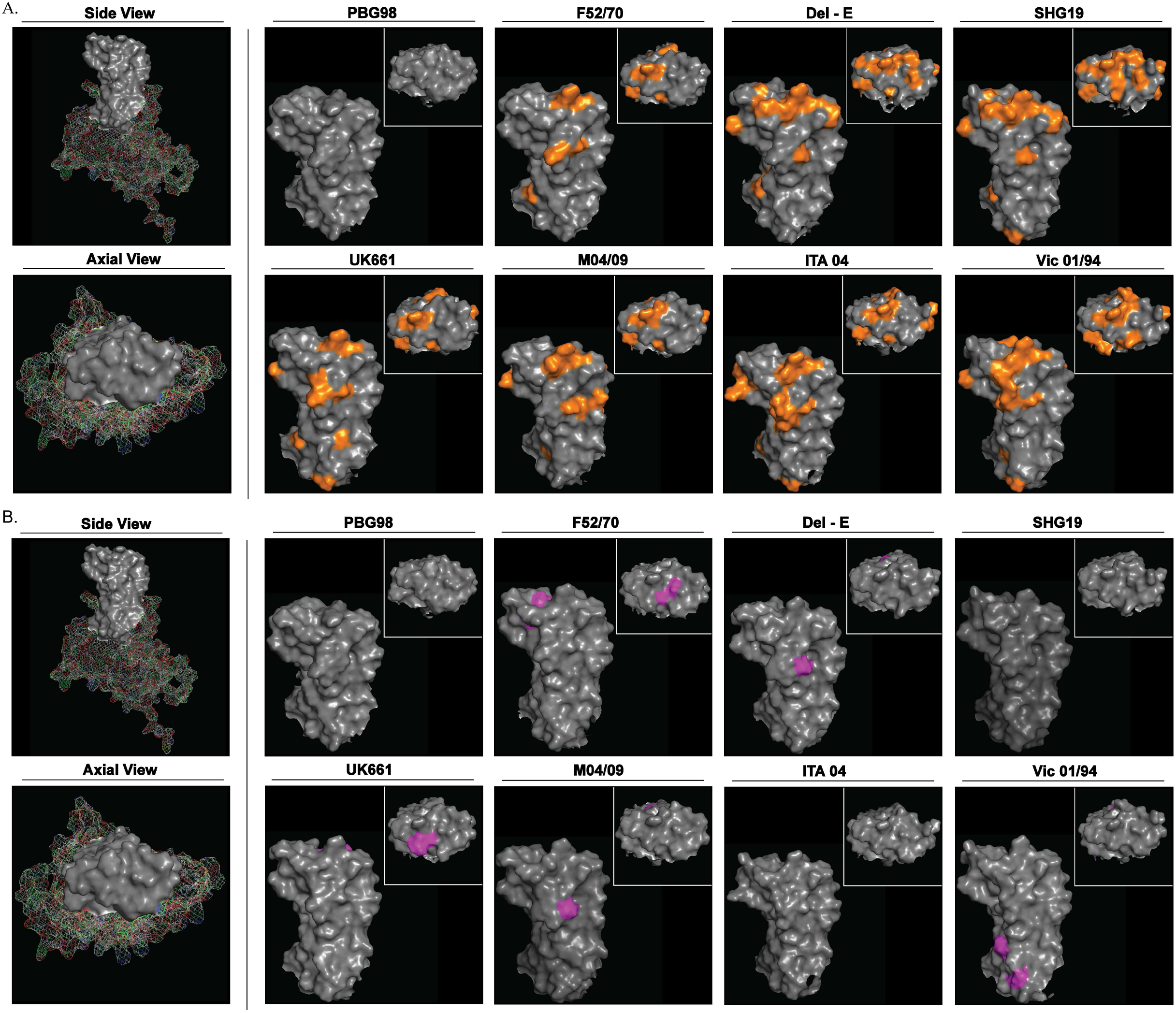
Structural modelling of the HVRs. The predicted structure of the VP2 of IBDV strain PBG98 was modelled using AlphaFold, and the side view and axial view were displayed. The HVR was depicted as solid grey. The predicted structures of PBG98, F52-70, Del-E, SHG19, UK661, M04/09, ITA-04, and Vic-01/94 were modelled using AlphaFold, based on the sequence that were in GenBank. The structures were compared to the PBG98 HVR structure and amino acid differences highlighted in orange (A). The predicted structures of the HVRs of the DT40-passaged viruses F52-70, Del-E, SHG19, UK661, M04/09, ITA-04, and Vic-01/94 were modelled using AlphaFold, and residues that were different from the GenBank sequences were highlighted in purple (B).

### Evaluation of the cross reactivity of serum neutralising antibodies from IBDV inoculated and vaccinated birds against the panel of diverse IBDV strains

Chickens were either inoculated with classical IBDV strain F52-70, or vaccine strain 228E (both genogroup A1). Twenty-eight days post inoculation, birds were humanely culled, bled, and the titer of serum neutralising antibodies determined against the panel of chimeric viruses (Figure 5). The virus neutralisation (VN) titer of antibodies from F52-70 inoculated birds against the F52-70 wt virus and the PBG98/HVR^F52-70^ chimeric virus (homologous controls) was 16.16 ± 0.29 and 15.41 ± 0.72, respectively, and there was no significant difference between them (p = 0.99), demonstrating that the chimeric virus was an adequate surrogate for the wt strain. There was also no significant difference in the VN titer between the PBG98/HVR^F52-70^ virus (genogroup A1) and the PBG98/HVR^UK661^ virus (A3) (13.32 ± 0.75, p = 0.664), or the PBG98/HVR^M04/09^ virus (A4) (11.41 ± 1.51, p = 0.063), whereas the PBG98/HVR^ITA-04^ virus (A6) and the PBG98/HVR^Vic 01/94^ virus (A8) were significantly less neutralised (10.41 ± 2.43, p = 0.0127; and 9.49 ± 1.66, p = 0.0029, respectively). The chimeric-viruses PBG98/HVR^DEL-E^ and PBG98/HVR^SHG19^ (both A2) were the least neutralized (7.91 ± 2.13, p = 0.0002, and 7.49 ± 0.76, p = 0.0001, respectively) (Figure 5A). The same pattern of neutralisation was observed with serum from 228E inoculated birds, where there was no significant difference in the VN titer against genogroup A1 viruses PBG98/HVR^F52-70^ or 228E wt (12.66 ± 1.38, and 12.82 ± 1.32, respectively, p > 0.99) whereas the genogroup A2 viruses, PBG98/HVR^DEL-E^ and PBG98/HVR^SHG19^, were significantly less neutralized (7.41 ± 2.02, and 7.41 ± 2.10, p = 0.0274). The titer against PBG98/HVR^UK661^, PBG98/HVR^M04/09^, PBG98/HVR^ITA-04^, and PBG98/HVR^Vic 01/94^ was 12.16 ± 1.26, 11.41 ± 1.28, 10.82 ± 0.90, and 9.32 ± 2.65, respectively, but there was no significant difference between them (Figure 5B).

**Figure 5.**
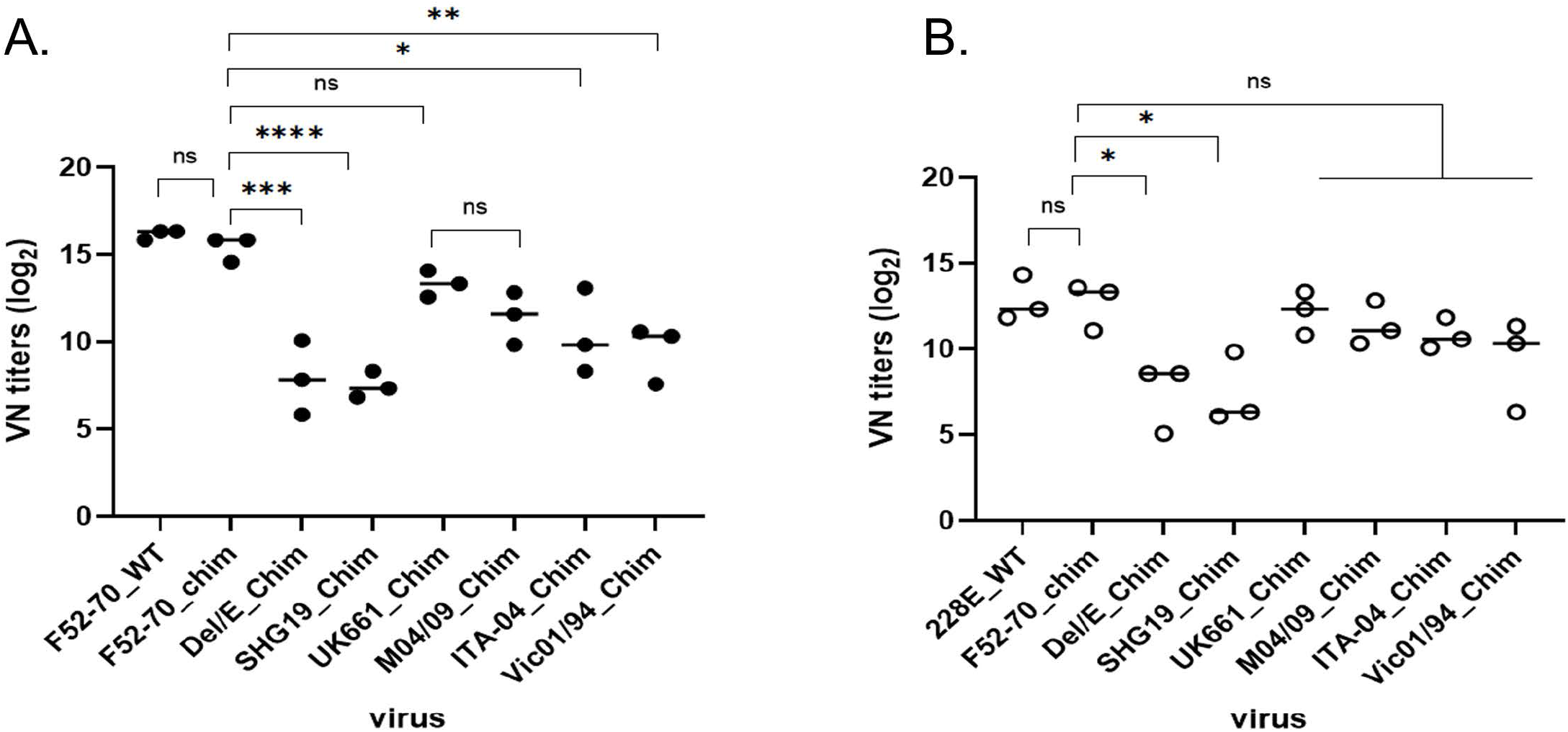
Quantification of the breadth of serum neutralizing antibody responses elicited by F52-70 and 228E against the panel. Viral neutralization assays were performed in DT40 cells, and the highest dilution of serum where there were no IBDV VP3 antigen positive cells was considered as the VN titer, which was expressed as log_2_ and plotted on a graph for birds inoculated with F52-70, closed circles (A) or 228E, open circles (B). The VN titer of the serum from three birds inoculated with F5270 and three inoculated with 228E was determined against chimeric-viruses containing the HVR from F52-70, Del-E, SHG19, UK661, M04/09, ITA-04, and Vic-01/94 and the backbone from PBG98 (*p<0.05, **p<0.01, ***p<0.001, ****p<0.0001).

## Discussion

The main correlate of protection described for IBDV vaccines is the neutralising antibody response against the VP2 capsid (35, 36). Worldwide, IBDV strains have been classified into 8 genogroups (A1-8), based on the sequence diversity of the VP2 HVR (9), however, the majority of traditional commercial vaccines have relied on a limited number of strains from genogroup A1, with little genetic diversity (36–39). Vaccination failures are being increasingly described in the field that are associated with mutations in the HVR, and there is a need for a method to screen traditional and contemporary vaccine candidates, platforms, and regimens for the breadth of immunity they elicit against different IBDV strains. Moreover, a method that could be used to identify immunodominant epitopes and conserved epitopes that induce more broadly cross-protective immune responses would be useful in informing the design of future vaccines. To address this, we developed a novel system for rescuing chimeric IBDVs and conducting neutralization assays, using the chicken B-cell line DT40. First, we demonstrated that DT40 cells could be used to titrate IBDV field strains of IBDV and conduct neutralisation assays. While DT40 cells have been previously shown to support the replication of IBDV (24, 25), field strains of IBDV continue to be titrated *in vivo*, by chicken infectious dose-50 (CID50) (40, 41), or *in ovo*, in embryonated eggs, by EID_50_ (19). Moreover, neutralisation assays continue to be performed either with embryonated eggs (42), or with cell-culture adapted strains of IBDV in primary chicken embryonic fibroblasts (CEFs) or the immortalised chicken embryonic fibroblast cell line DF-1 (41, 43). Here we demonstrated that there was a linear relationship between TCID_50_ determined in DT40 cells, and EID_50_ (R^2^ = 0.9313), providing support for using DT40 cell TCID_50_ as a surrogate of EID_50_. Moreover, we demonstrated that the cells can also be used to quantify the titer of neutralising antibodies against field strains.

Next, we demonstrated DT40 cells could be used to rescue a molecular clone of IBDV. Traditionally, recombinant strains of IBDV have been rescued by transfecting DF-1 cells with plasmids encoding segments A and B. While this reverse genetics system is efficient, it can only be applied to cell-culture adapted strains of IBDV that replicate within DF-1 cells. Recently, it has been shown that the cell lystates from transfected DF-1 cells can be passaged onto chicken primary bursal cells to rescue a molecular clone of a field strain, which does not replicate within DF-1 cells (26). Here, we extend these observations by electroporating DT40 cells with plasmids encoding IBDV segments A and B, to rescue a molecular clone of IBDV in the absence of DF-1 cells. We then used this system to rescue recombinant chimeric IBDVs containing the HVR from seven diverse field strains from six different genogroups. Recombinant viruses replicated to high titers in our system, demonstrating that this approach was efficient. To date, comparative antigenicity studies have been limited to laboratories with access to diverse IBDV field strains, and the rescue of IBDV field strains has been limited to labs with *in vivo* facilities to provide a supply of primary B cells, however, our approach can enable studies to be conducted in a wider number of labs, as DT40 cells are immortal and commercially available, and the rescue system can be applied to any strain where the sequence is known.

We then determined the ability of serum from IBDV-inoculated birds to neutralize the panel of chimeric viruses. Based on our neutralization data, sera from chickens inoculated with genogroup A1 viruses F52-70 or 228E had the lowest VN titers against genogroup A2 viruses PBG98/HVR^Del-E^ and PBG98/HVR^SHG19^. These data are consistent with what is observed in the field, where the emergence of genogroup A2 viruses in the 1980s in the USA necessitated the use of alternative vaccines, as traditional live vaccines based on A1 strains only partially protected flocks (44, 45). Variant strains belonging to genogroup A2 have also more recently emerged in China and are not adequately controlled by vaccines against vv strains (15, 46). In contrast, there was no significant difference in the VN titer of serum from birds inoculated with F52-70 or 228E against the homologous A1 strain (PBG98/HVR^F52-70^), the A3 strain (PBG98/HVR^UK661^), or the A4 strain (PBG98/HVR^M04/09^), suggesting that these genogroups are antigenically more closely related to genogroup A1 strains, and that vaccines containing the VP2 from genogroup A1 strains may be likely to provide better protection. These data are also consistent with what is observed in the field, where vaccines including the VP2 from genogroup A1 viruses are used to control vv IBDV strains belonging to genogroup A3 (36), and distinct A4 strains in South America (47).

Using serum from F52-70 inoculated birds, we observed that the genogroup A6 and A8 viruses (PBG98/HVR^ITA-04^ and PBG98/HVR^Vic-01/94^) were significantly less neutralised than the A1 strain (p < 0.05 and p < 0.005, respectively), suggesting that they are antigenically more distant from genogroup A1 viruses and that vaccines including the VP2 antigen from A1 viruses may be less efficacious. Consistent with this observation, it was shown that vaccines based on A1 strains were not protective against the ITA-04 (genogroup A6) strain (48), and the majority of Australian IBDV strains are controlled through the use of genogroup A7 vaccines such as V877 and 002/73 (9, 49), and not through the use of genogroup A1 vaccines, although Vic-01/94, included in our panel, is variant strain belonging to genogroup A8, and outbreaks have been reported in vaccinated flocks (50). Using serum from 228E inoculated birds, we observed that the VN titers against genogroup A6 and A8 viruses were not significantly different from those of A1 strains, however, given that we observed the same trend in the data with birds inoculated with F52-70, these data may reach statistical significance if more birds were used per group.

Taken together, our neutralisation data are consistent with observations from the field, validating the proof of concept that we can determine the neutralization profile of vaccine serum against diverse IBDV field strains using our novel chicken B-cell rescue system and neutralisation method. Based on our observations, genogroup A3 and A4 are likely to be more closely antigenically related to genogroup A1 strains than genogroups A6 and A8, and genogroup A2 is likely to be the most antigenically distant from genogroup A1 strains. It should be noted that commercial HVT-based vectored vaccines containing theVP2 gene from the genogroup A1 strain F52-70 do show protection against the US variant Del-E (genogroup A2) strain (39). However, more studies are needed to understand the relative contribution of T cell mediated immunity and neutralizing antibody responses to the protection provided by HVT-based vaccines. Moreover, while present study was concerned with antibody responses elicited by live viruses, our method could be also applied in the future to compare the breadth of antibody responses elicited by different vaccine platforms, including HVT-based vaccines (37, 39).

Following five passages in DT40 cells, two of our recombinant chimeric viruses (PBG98/HVR^SHG19^ and PBG98/HVR^ITA-04^) had no mutations in the HVR, but four viruses (PBG98/HVR^F52-70^, PBG98/HVR^Del-E^, PBG98/HVR^UK661^, and PBG98/HVR^M04/09^) developed two mutations: S251I and S315Y (F52-70), C262Y and N279H (Del-E), T250S and S251I (UK661), and V256L and N279H (M04/09), and one virus (PBG98/HVR^Vic 01/94^) developed six mutations (T227I, T272I, S299N, E324Q, L328S, V329A) (Figure 4). It remains unknown why the virus carrying the Vic 01/94 HVR had more mutations than the other strains. Neutralization assays are typically conducted with cell-culture adapted viruses that usually contain mutations in the HVR, and it is challenging for neutralizing antibodies to be characterized against field-strains with wild-type HVR sequences. Adapting IBDV field strains to replicate in immortalized adherent cell-culture is associated with mutations at amino acid positions 253, 279, 284 and 330 in the HVR, which are known to change antigenicity and virulence (12–14, 26, 34). However, we demonstrated that there was no change in the residues at positions 253, 284 and 330 in our recombinant viruses, although strains Del-E and M04/09 had an N279H mutation following DT40 passage. In the past, it has also been shown that amino acid substitutions at position 279 are involved in DT40-cell adaptation: the classical virulent strain, GBF1, developed an N279Y/H mutation, and the lab-adapted strain, Soroa, developed an N279D mutation (24, 25), but the vv strain, Lx, did not (24, 51). We also detected mutations S315Y in strain F52-70, and V256L in strain M04/09 following DT40 cell passage, and it has previously been reported that the DT40 cell-adapted IBDV strain Soroa had mutations at the same positions (S315F and V256A) (25). The T250S mutation we observed in UK661 has also been reported in the genogroup A7 Australian strain 002-73 by a phage display method. While the mutation was associated with reduced binding of monoclonal antibodies to a conformational epitope, it has not been further characterized in cell culture experiments with other IBDV strains (52). Of the mutations we observed in Vic 01/94, the presence of Isoleucine (I) at position 272 is suspected to be associated with virulence, and threonine (T) with attenuation (53, 54), although neither had an effect on neutralizing epitopes (53), the S299N mutation has also been observed in classical virulent F52-70 and antigenic variant Del-E strains, and the E324Q, L328S and V329A mutations are reported to be part of the “QMSWSASGS” signature of virulence (residues 324 to 332) (16, 55), but have not been associated with changes in antigenicity. To our knowledge, the other mutations we observed (S251I, C262Y, and T227I) have not been previously described. Interestingly, the C262Y, and the T227I mutations changed the residue to the amino acid that is present in all other strains in the panel, and the S251I mutation arose both in the A1 virus F52-70, and the A3 virus, UK661. Taken together, some of our rescued strains had mutations consistent with DT40 cell-adaptation, however, whether these mutations altered IBDV antigenicity remains to be determined, and, given that the pattern of neutralisation we observed was consistent with field observations, we believe that our neutralization data are still relevant to the field.

When we modelled the structure of the HVR based on the GenBank sequences, the viruses belonging to genogroup A2 (Del E and SHG19) had more extensive changes to the axial view of the HVR compared to the backbone (PBG98, genogroup A1) than strains belonging to the other genogroups F52-70 (A1), UK661 (A3), M04/09 (A4), ITA-04 (A6), and Vic 01/94 (A8). This is consistent with them having the lowest VN titers, demonstrating that the predicted HVR structures correlated with the patterns of antigenicity, thus linking the IBDV HVR sequence, structure, and antigenicity. When we compared the structures modelled from the GenBank sequences to the structures modelled from the passaged viruses, we found that only the DT40 cell adaptation mutations S251I and S315Y in F52-70, and T250S and S251I in UK661 were on the axial view, whereas the other mutations, including position 279, and the six mutations in Vic-01/94, were located on the side of the VP2 molecule, suggesting that IBDV may not rely solely on the axial tip of the VP2 for binding the receptor on DT40 cells. Defining which residues are involved in binding the canonical receptor on chicken B cells *in vivo* is an important question to address in the future.

In summary, we have developed a novel IBDV rescue system and neuralization assay using the chicken B-cell line, DT40. We used this method to engineer a panel of seven recombinant viruses containing the HVR from six different genogroups, and we characterised the breadth of neutralizing antibodies generated by genogroup A1 strains F52-70 and 228E against the panel. Our data are consistent with field observations, validating our approach, and we can use our method in the future to screen novel IBDV vaccine candidates, platforms, and regimens for cross reactivity against different genogroups. In addition, we will be able to perform cross-neutralization studies to evaluate the antigenic relatedness of diverse field strains, providing valuable information on how sequence diversity relates to antigenic diversity that could inform vaccine design in the future. Moreover, coupling our approach with protein modelling, we will be able to determine the contribution individual amino acids make to antigenicity, and define immunodominant and conserved epitopes for the rational design of future vaccines.

## Acknowledgements

This work was supported by grants BBS/E/I/00001845, BBS/E/I00007034, and BBS/E/I/00007039, funded by Biotechnology and Biological Sciences Research Council, U.K. The funders had no role in study design, data collection and interpretation, or the decision to submit the work for publication.

## Notes

### Competing Interest Statement

The authors have declared no competing interest.

